# Global overview of sea turtle fibropapillomatosis tumors: a survey of expert opinions and trends

**DOI:** 10.1101/2024.06.06.597728

**Authors:** Jenny Whilde, Narges Mashkour, Samantha A. Koda, Catherine B. Eastman, Drew Thompson, Brooke Burkhalter, Hilary Frandsen, Annie Page, Nicholas B. Blackburn, Karina Jones, Ellen Ariel, Sophie M. Dupont, Lawrence Wood, David J. Duffy

**Author notes:** Corresponding author: David J. Duffy. These authors contributed equally to this work (Jenny Whilde, Narges Mashkour).

## Abstract

Marine environments offer a wealth of opportunities to improve understanding and treatment options for cancers, through insights into a range of fields from drug discovery to mechanistic insights. By applying One Health principles the knowledge obtained can benefit both human and animal populations, including marine species suffering from cancer. One such species is green sea turtles (*Chelonia mydas*), which are under threat from fibropapillomatosis (FP), an epizootic tumor disease (animal epidemic) that continues to spread and increase in prevalence globally. In order to effectively address this epizootic, a more thorough understanding is required of the prevalence of the disease and the approaches to treating afflicted turtles. To identify knowledge gaps and assess future needs, we conducted a survey of sea turtle FP experts. The survey consisted of 47 questions designed to assess general perceptions of FP, the areas where more information is needed, local FP trends, the disease status, and mitigation needs, and was voluntarily completed by 44 experts across a broad geographic range. The survey responses provided a valuable overview of the current FP status in sea turtles, FP research, and insight into the approaches currently taken by turtle rehabilitation facilities around the world. Over 70% of respondents both recognized FP as a cancerous panzootic disease, and reported that FP is increasing in prevalence. They report several factors contributing to this increase. Nearly all of the respondents reported that FP research, patient treatment and rehabilitation required more funding in their area, and reported inadequate facilities and capacity for dealing with FP patients. Treatment approaches varied: just over 70% of the medical experts that responded surgically remove FP tumors, either using laser or scalpel. Just under half of respondents use anti-cancer drugs in their treatment of FP. Internal tumors were reported as justification for euthanasia by 61.5% of respondents, and 30.8% reported severe external tumors to be sufficient grounds for euthanasia. Most medical respondents (93.3%) routinely perform necropsy on deceased or euthanized FP-afflicted turtles. Over 80% of respondents considered large-scale multidisciplinary collaboration ‘extremely important’ for advancing the field of FP research. The survey responses provide a valuable insight into the current state of FP treatment, rehabilitation and research, and help to identify critical FP-related research and rehabilitation areas most in need of attention.

## Introduction

The ocean can provide novel anti-cancer therapeutics, insights into naturally occurring cancers, and understanding of the complex interplay between environmental and viral triggers of tumor development^1,2^. Taking a One Health perspective can simultaneously increase our understanding of and ability to treat both human and animal cancers ^3-5^. Sea turtles are long-lived vertebrates exposed to multiple oceanic habitats, so they offer valuable insights into marine exposures and mechanisms promoting tumorigenesis^2,6^. Sea turtles are categorized by the IUCN on a spectrum of vulnerable to critically endangered^7^, and are subject to multiple anthropogenic and environmental pressures^8-14^. Moreover, in some regions there is an extra challenge to their survival: the neoplastic disease fibropapillomatosis (FP). FP is a debilitating disease that afflicts all seven species of sea turtles^15^, with a higher prevalence in green turtles (*Chelonia mydas*)^16^. The disease has reached epizootic status in some populations of green turtles (e.g. Florida, USA)^17^ and continues to spread to regions where it has not been reported before^18-24^.

Fibroepithelial FP tumors are formed on the soft external tissues and occasionally on the carapaces and plastrons (upper and lower shell) of sea turtles. External tumors can vary in size and distribution (Fig. 1a). Higher burdens of tumors can significantly impede locomotion, foraging ability, vision, and predator evasion^15^. When they occur internally, fibroma FP tumors affect the vital functions and survival of sea turtles. Internal tumors are frequently observed in lungs, kidneys, and livers, and, to a lesser extent, bladders, mouths and bones^25-28^.

**Figure 1.**
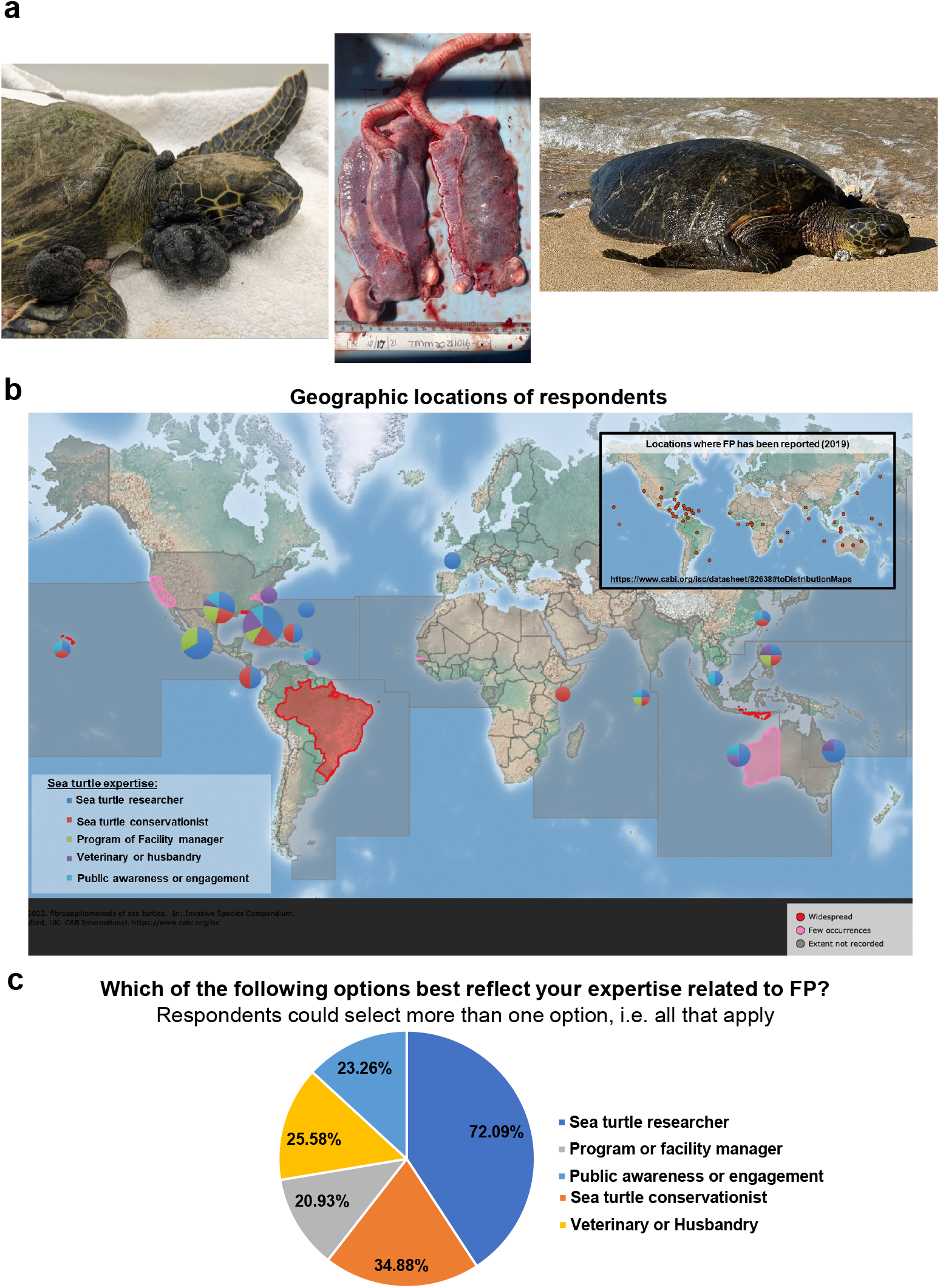
**a) Left**: FP-afflicted green sea turtle patient at the University of Florida Whitney Laboratory for Marine Bioscience and Sea Turtle Hospital. Patient stranded with monofilament entanglement and severe FP tumor burden, including advanced eye tumors. **Center**: The lungs of a deceased green sea turtle patient at the UF Whitney Lab Sea Turtle Hospital, with multiple internal tumors present in both lobes. **Right**: FP-afflicted sea turtle on a beach in Maui, Hawai’i. Image: Matt Belonick, 2022. **b)** Global map of the geographic location of respondents. Pie charts indicate the location and sea turtle expertise area of respondents. Chart sizes are proportional in size to the number of respondents from that location. Inset map: locations where sea turtle FP has been reported, data from CABI: https://www.cabi.org/isc/datasheet/82638#toDistributionMaps. CABI map is licensed under a Creative Commons Attribution-NonCommercial-ShareAlike 3.0 Unported License. **b)** Pie chart showing the self-reported sea turtle expertise area of each respondent.

Juvenile turtles are at higher risk of developing FP when they enter neritic foraging areas after completing their pelagic life stage^29^, due to the infectious nature of the disease, dietary shifts associated with this life-stage change, and/or proximity to near-shore contaminants from anthropogenic activity. To date, at least one virus, chelonid alphaherpesvirus 5 (ChHV5), has been strongly associated with FP while other viruses such as *Chelonia mydas* papillomavirus 1 (CmPV1), retrovirus and sea turtle tornovirus 1 have been found in tumor tissues of green turtles with no clear association to disease development, pattern, and severity^30-40^. Disease manifestation is likely to be multifactorial, as the presence of ChHV5 alone is not linked with tumor formation, and recovery and regression are possible in some cases^25,41-45^. Anthropogenic stressors, environmental pollutants, immunosuppression and genetic predisposition are suggested as co-factors in tumor growth and disease prevalence^15,26^. Researchers have focused on various aspects of FP, from causal agents and prevention to rehabilitation and population dynamics after release^15,16,20,24,30,37,46,47^. The use of novel oncology techniques has also been reported in combatting FP^41,42,48-50^.

Overall, the complexity of the disease puts pressure on conservation efforts. In some areas, stranded turtles are treated with laser surgery to excise external tumors. If the tumor burden is high, or internal tumors are detected, the animal will be euthanized. Currently, euthanasia is commonly utilized for sea turtles with internal tumors. The hard outer shell of sea turtles prevents surgical access, and, as no other treatments exist yet, internal tumors are generally inoperable. There is no universal international policy for determining when to operate on turtles with FP, and decisions are made by local veterinarians on a case-by-case basis depending on the FP burden of the turtle. Existing FP tumor scoring systems have been suggested to be utilized to aid treatment decisions^51,52^. As the disease continues to emerge in new regions, questions of risk and management remain largely unanswered.

In October 2021, a two-day international symposium on sea turtle FP research was hosted virtually via Zoom by the University of Florida Whitney Laboratory for Marine Bioscience and Sea Turtle Hospital in St. Augustine, Florida. In total, 148 researchers, veterinarians, rehabilitation facility managers, conservationists and stakeholders attended the symposium over the two-day period. Twenty experts presented their findings orally and shared their knowledge with the audience on different aspects of FP in sea turtles. To further inform the scientific community we conducted an FP-focused online survey which was voluntarily completed by symposium attendees who self-identified as FP experts. The responses to the survey questions were analyzed and are reported here in hopes of identifying FP trends and defining the critical FP-related research and rehabilitation areas most in need of attention.

## Methods

A total of 47 questions were formulated to assess general perceptions of FP and the areas where more information is needed. The questionnaire was designed based on published literature and discussions with experts in the field. The questions encompassed current local FP trends, the possibility of updating the disease status, and mitigation needs. Sixteen questions were designed exclusively for veterinarians, veterinary technicians and assistants, husbandry staff, and rehabilitation managers to collect information on the various approaches to FP treatment.

The questions were presented online via the Survey Monkey platform (www.surveymonkey.com) and the symposium attendees received Mailchimp notifications with a link to respond to the questions after they consented to participate. Attendees were informed of the project goals and the participation conditions during the symposium via both oral and written communication.

Responses were collected anonymously; however, to be informed of regional opinions and decisions, the respondents were asked to voluntarily identify their countries and regions. Their area of expertise or their field was also requested. When possible, the questions were open-ended to avoid funneling responses towards a particular answer. To deter any bias introduced by the question format, a comment section was provided in case the given numerical values or multiple-choice answers to the questions were not suitable or applicable. Respondents could also leave answers blank if they were not applicable.

The first section of the survey aimed to obtain information on the current incidences of FP and the treatment, management, and conservation options that are implemented in the respondents’ areas. Respondents were asked to define the species present in their area and whether they had encountered turtles afflicted with FP. They were then asked to rate the known threats to sea turtles and to specifically rank the severity of the threat of FP to sea turtle conservation. The co-factors of FP which have been reported or suggested in the scientific FP literature were presented to the respondents to investigate their opinion of the relevance of these co-factors. The second survey section focused on research priorities and the impediments to achieving important FP research goals. The third section of the survey gathered information on FP treatments, rehabilitation, and research activities that are currently taking place in respondents’ facilities or institutions in their regions. Finally, the fourth section was available exclusively for veterinary, rehabilitation, rescue and husbandry experts, as this section focused on respondents’ opinions on rehabilitation and treatment procedures. Respondents with medical expertise were asked questions about their rehabilitation facilities, treatment plans, equipment, recovery rates, and decision-making processes for releasing or euthanizing FP-afflicted turtles.

## Results Respondent profile

Of the 148 symposium participants, 44 experts voluntarily participated in this survey. The response rate varied by question and ranged from 84% to 97%. Participation was circum-global, representing the majority of the locations where FP has been reported in sea turtles (Fig. 1b). Respondents categorized themselves as researchers (72.1%), conservationists (34.9%), program or facility managers (20.9%), veterinarians and husbandry staff (25.6%) and public awareness or engagement professionals (23.3%) (Fig. 1c). Note that the total percentage is 176.8% as respondents could select all descriptions that applied to them (e.g., a respondent could be both a veterinarian and a researcher).

Twenty-four out of 44 respondents were from the USA, with 40% of respondents working in Florida, USA. This was expected, as the symposium was hosted in Florida, which is a hot spot for FP^15,26^. The first reported case of FP in the scientific literature was in the Florida Keys in 1938^53^, with anecdotal reports of FP occurring at low prevalence in Florida from the early 1800s ^54^. The most abundant species in the areas where the respondents work are green turtles, followed by loggerheads (*Caretta caretta*) and hawksbills (*Eretmochelys imbricata*) (Fig. 2a). Fibropapillomatosis was observed by respondents in all species of sea turtles except flatback turtles (*Natator depressus*), with 100% of respondents reporting observations of green turtles with FP tumors (Fig. 2b). Observations of FP in all seven sea turtle species have been reported in the scientific literature, but green turtles are the most frequently afflicted and reported^15,55^.

**Figure 2.**
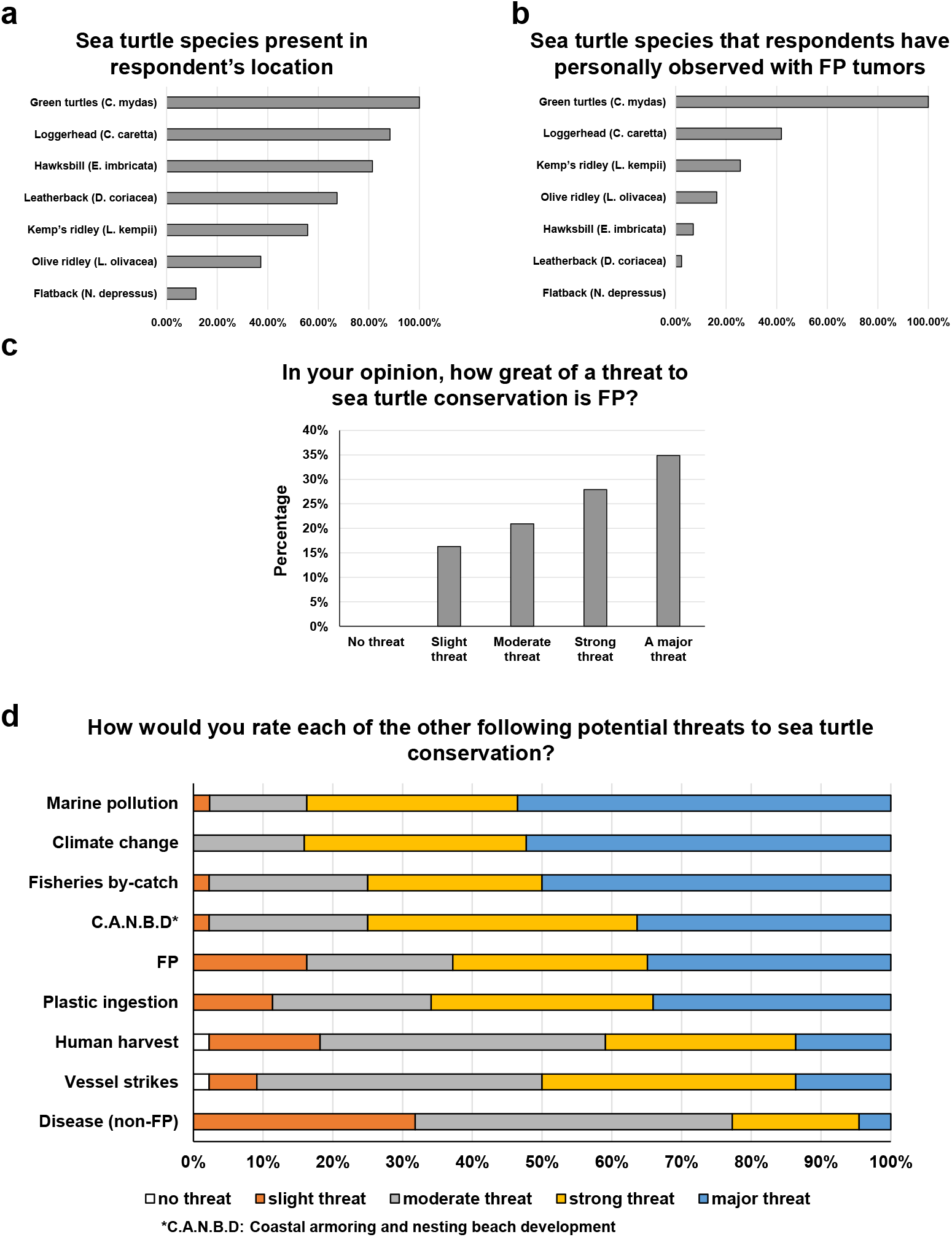
**a)** Percentage of respondents reporting the presence of each sea turtle species present in their location. **b)** Percentage of respondents that reported personally observing FP tumors in each sea turtle species. **c)** Percentage of respondents that consider FP to be no threat, slight threat, moderate threat, strong threat or a major threat to sea turtle conservation. **d)** Proportion of respondents that consider potential threats to sea turtle conservation to be no threat, slight threat, moderate threat, strong threat or a major threat.

### The threat of FP to sea turtle conservation

All expert respondents believed that FP poses a threat to sea turtle conservation, with 34.9% of respondents considering it to be a major threat (the highest threat level option) (Fig. 2c). Of the nine threats to sea turtle conservation that were assessed, three were identified as major threats by over 50% of respondents (Fig. 2d). These three threats in order of most severe were: Marine Pollution, Climate Change and Fisheries By-catch. A further three threats were considered to be major threats by over a third of respondents: Coastal Armoring and Nesting Beach Development, FP, and Plastic Ingestion (Fig. 2d). These threats are not mutually exclusive but can be interrelated or have additive, or even negatively synergistic effects. For example, climate change and marine pollution are believed to exacerbate human and wildlife disease, likely including FP which has environmental co-triggers.

Over 74% of expert respondents consider FP to be a panzootic (Fig. 3a) - an animal pandemic that occurs in a widespread outbreak among a large number of animals, usually affecting more than one species. Seventy-two percent of respondents reported FP to be increasing in their region, with only 5% reporting a decrease and 23% remaining steady (Fig. 3b). Reflecting the current literature, only green turtles were reported to have more than 25% FP prevalence in any region (Fig. 3c). Nearly 12% of respondents reported that 51-75% of the green turtles were afflicted by FP in their region, while 9.5% of respondents reported that over 76% of green turtles in their region were FP-afflicted (Fig. 3c). These estimates were primarily derived from Stranding Data, In-water Data and Personal Observation, with each respondent citing a mixture of data sources (Supplemental Fig. 1a). In the majority of regions, the prevalence of FP at the population level was reported as increasing (Fig 3b), as was the tumor burden experienced by each individual turtle (52.4%) (Fig. 3d). Only 2.4% of respondents reported a decrease in individual tumor burdens, while 45.2% reported static tumor burdens.

**Figure 3.**
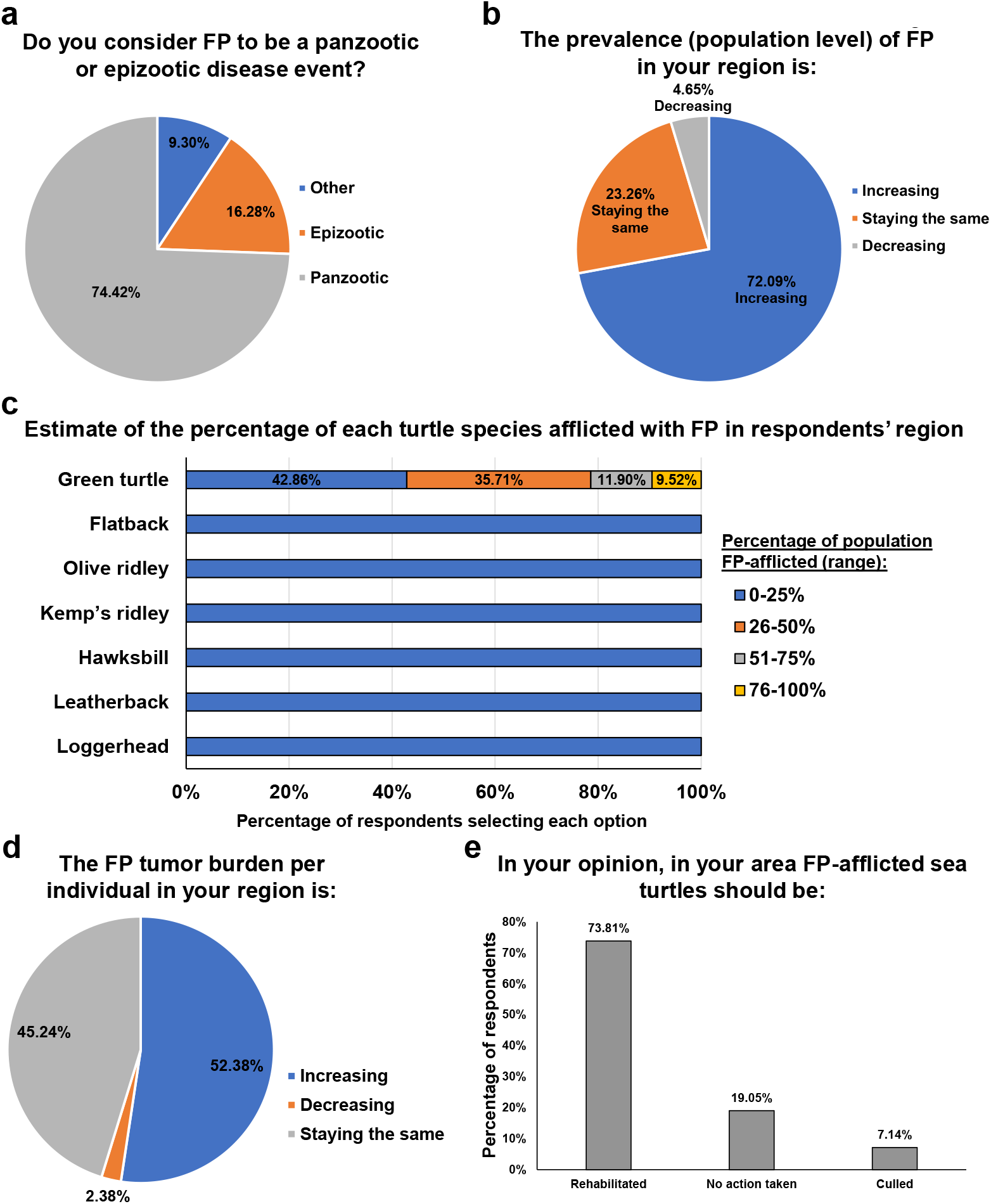
**a)** Percentage of respondents that consider FP to be a panzootic or epizootic disease event. **b)** Percentage of respondents that consider the prevalence of FP in their region to be increasing, staying the same, or decreasing. **c)** Estimated percentage of each sea turtle species afflicted with FP in the respondent’s region. **d)** Percentage of respondents that consider the FP tumor burden per individual in their area to be increasing, staying the same, or decreasing. **e)** Percentage of respondents that think that FP-afflicted sea turtle should be rehabilitated, culled or no action taken.

Given the general increasing trend of FP in green turtles, we next assessed regional differences in FP management. Over half (58.5%) of respondents reported that, in their region, rehabilitation of FP-afflicted sea turtles was carried out, 34.2% reported that currently no action is taken in relation to FP turtles, and 7.3% reported that sea turtles stranded with FP are culled. The course of action that respondents believed to be optimal for FP-afflicted sea turtles closely mirrored the reported strategies currently in place in their region: 73.8% of respondents believed rehabilitation should be conducted, 19.1% believed that no action was necessary, and 7.1% believed animals should be culled (Fig. 3e). While rehabilitation was reported as the most common response to FP, a further 36.9% of experts in regions where rehabilitation is not conducted (an additional 15% of total respondents) believe that rehabilitation should be applied.

### Causative factors of FP

The next set of questions focused on canvasing respondent’s opinions on the nature of FP and its causative factors. The majority of respondents (70%) consider FP to be a cancerous disease (Fig. 4a). Much of the historical FP literature mainly refers to FP as a tumor disease, and prior to that as warts, but recent advances in FP genomics and transcriptomics have shown that FP shares gene expression changes and mutational profiles with human cancers, with sea turtle FP oncogenic signaling being strikingly similar to human pan-cancer oncogenic signalling^17,26,41,42^. The respondents’ understanding of FP reflects these recent advances in FP research.

**Figure 4.**
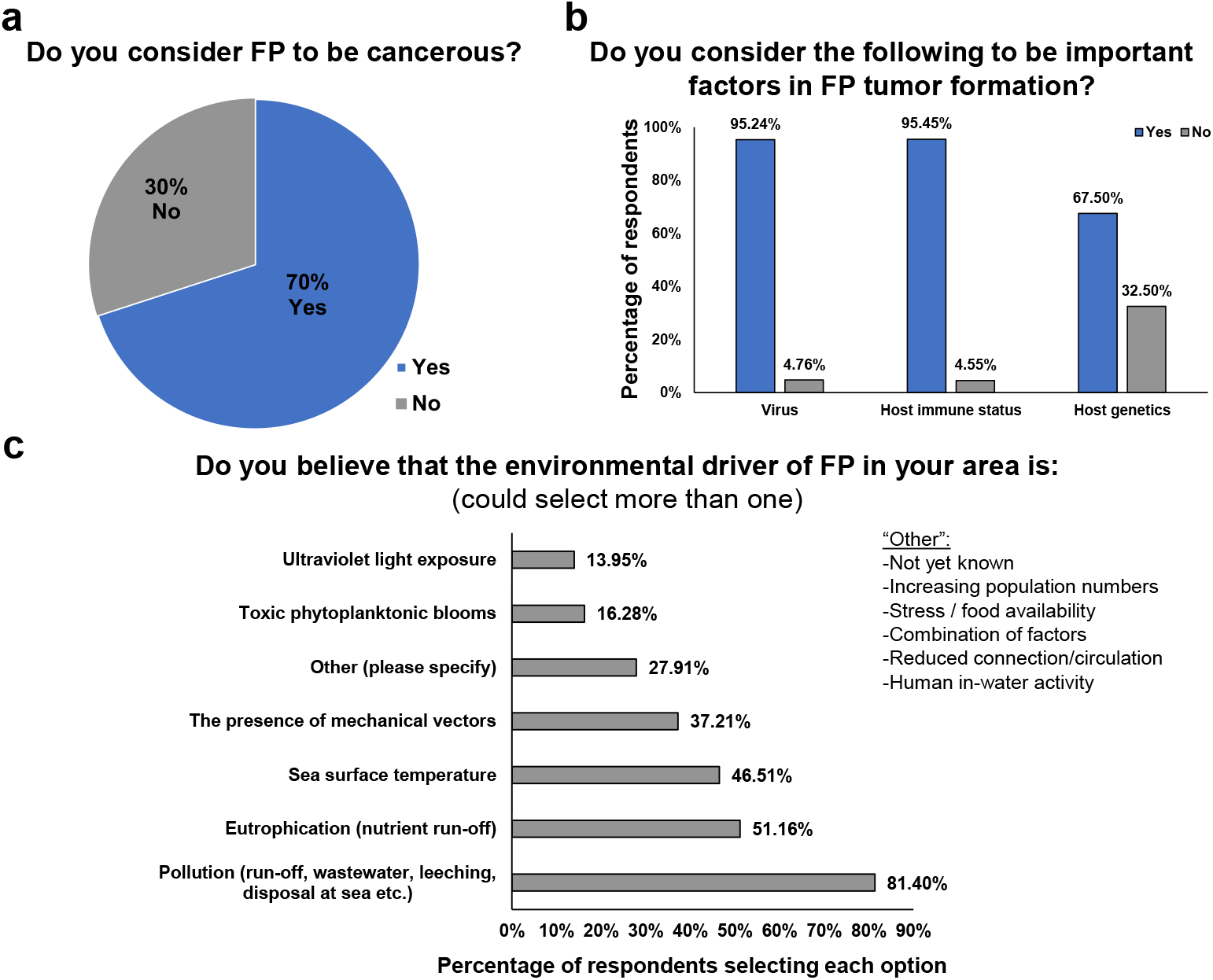
**a)** Percentage of respondents that consider FP to be cancerous or non-cancerous. **b)** Percentage of respondents that consider viruses, host immune status and host genetics to be important factors in FP tumor formation. **c)** Factors that respondents believe are environmental drivers of FP in their area.

Respondents consider viral (95.2%), host immune status (95.5%) and host genetic factors (67.5%) to be important for FP tumor formation (Fig. 4b). The majority of respondents (81.4%) believe that pollution is the environmental driver of FP in their area, followed by eutrophication (nutrient run-off, 51.1%) (Fig. 4c). Interestingly, all of the available options were selected by one or more respondents, highlighting the current lack of definitive causal evidence and the multi-factorial nature of the disease’s environmental cofactors (Fig. 4c).

### FP research, rehabilitation and management: funding status and future priorities

For 75.9% of respondents, the largest impediment to conducting FP-related research is a lack of available funding. Regulatory requirements, permits, low recapture rates, infrastructure and training were also cited as impediments. When queried directly whether FP research in their area received adequate funding, 94.9% of experts reported that more funding is required, 5.1% reported research funding levels to be adequate, and 0% reported that FP research should receive less funding. Nearly half of respondents (48.7%) reported that governmental support was the primary source of FP funding in their area (Fig. 5a).

**Figure 5.**
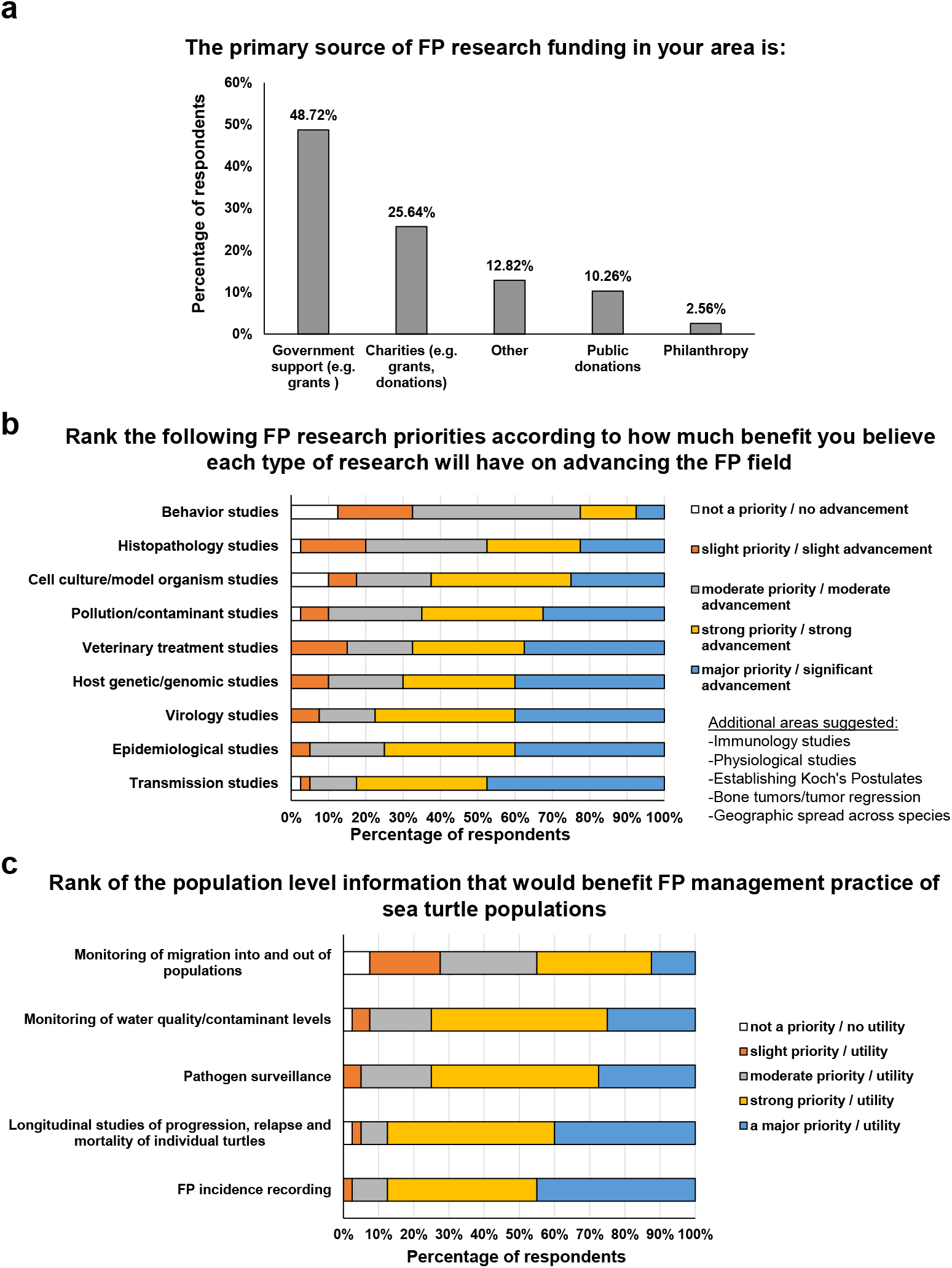
**a)** Percentage of respondents that receive FP funding research from government support, charities, public donations, philanthropy or other source. **b)** Amount of benefit that respondents believe each type of research will have on advancing the FP field. **c)** Amount of benefit that each type of population-level information is considered to be for FP management practices.

For rehabilitation, funding was reported as an impediment by only 21% of responders. Instead, a lack of facilities (non-existent or inadequate capacity) or limited expertise was cited as an impediment by 69.6% of respondents. However, when asked directly, 79.0% of expert respondents reported that FP rehabilitation and treatment in their area required more funding, with 15.8% reporting that it was adequately funded and 5.3% reporting that it should receive less funding. The primary defined source of rehabilitation funding is Philanthropy (26.3%), followed closely by Public Donations (23.7%), and 15.8% reporting Governmental sources as their primary funder (Supplemental Fig. 1b). The remaining 34.2% of respondents selected Other, but over half of this number constituted respondents that reported no funding availability in their area.

In the majority of respondents’ areas (78.4%) there is inadequate capacity for FP rehabilitation and rescue. Only 13.5% of respondents reported adequate capacity, while 8.1% reported their capacity as being too high. On average, facilities can accommodate and care for an average of nine turtles at any one time (range = 0-50). A respondent from the Philippines reported that no dedicated sea turtle medical facility exists in their area and that rehabilitation is currently performed in private homes, in government offices or gardens, or at dive centers, and expressed concern about the inability to have rigorous biosecurity or safe treatment under these conditions. Another respondent reported zero capacity at dedicated facilities, instead working in conjunction with local veterinary clinics. In general, 67% of sea turtle rehabilitation/rescue facilities reported that their capacity was too low, 24.6% of respondents reported their general sea turtle facilities had adequate capacity and 7.7% reported that their facility capacity was too high.

Respondents cited insufficient knowledge as the main impediment to implementing FP-related population management strategies. This included basic knowledge about the disease and its spread, and a lack of knowledge on the overall FP status of local populations. The majority of respondents believe that all research priority areas they were asked about would be beneficial and advance the FP field of study (Fig. 5b). Of all research areas listed, transmission studies were ranked as the area that should be a major priority as it would provide significant advancement (47.5% of respondents assigned this the highest priority ranking, Fig. 5b). Epidemiological, virology and host genetic/genomic studies all tied second place in terms of ranked importance to the field of FP research (40% of respondents assigned these the highest priority ranking, Fig. 5b). Respondents considered ‘FP incidence reporting’ would have the highest utility for population-level FP management practice (45% of respondents assigned this the highest priority ranking, Fig. 5c), followed closely by ‘Longitudinal studies of progression, relapse and mortality of individual turtles’ (40%, Fig. 5c).

### Clinical care of FP-afflicted sea turtles

The majority of the 14 respondents with medical expertise (71.4%) reported that FP tumors are surgically removed in their location, with 28.6% reporting no surgery is conducted. Of those that conduct surgery, 70% excise FP tumors by laser surgery, while 30% of respondents use a scalpel. Three drugs were reported as being in use for FP treatment: acyclovir (anti-viral), 5-fluorouracil (5-FU, anti-cancer) and bleomycin (anti-cancer), each of which has been reported in the literature^41,48-50^. Over half of respondents do not use anti-cancer drugs in their treatment of FP (57.1%), while 42.9% do use them.

The criteria for making a euthanasia decision varied between respondent regions. Internal tumors were reported as justification for euthanasia by 61.5% of respondents, 21.3% cited the presence of ocular tumors, and 15.4% said that euthanasia would only be justified in the case of bilateral visual defects. Nearly one-third (30.8%) reported severe external tumors as being sufficient grounds for euthanasia, while in more remote locations in Western Australia it was reported that FP-afflicted animals are euthanized if no rehabilitation facilities are available. One respondent reported that euthanasia of FP-afflicted turtles is never authorized in their country because they are a protected wildlife species. Another respondent reported that in their region FP-afflicted turtles are not euthanized but are released after the maximum amount of veterinary care available has been provided. Most medical respondents (93.3%) routinely perform necropsy on deceased or euthanized FP-afflicted turtles.

Over seventy percent (71.4%) of respondents had access to CT scans and x-rays for their patients (either at their facility or at a provider they could use), while 28.6% did not. The number of respondents with access to CT scans closely mirrored the number of respondents that assess for internal tumors; 78.6% of medical respondents assess for internal tumors, while 21.4% do not. Post-treatment FP regrowth rates at the facilities holding sea turtles after treatment is unknown in 50% of cases, while 21.4% of respondents reported regrowth rates in 0 - 25% of patients (Fig. 6a). An equal number of respondents (14.3% in both) reported that 26-50% of patients exhibit regrowth and 51% -75% of patients had regrowth (Fig. 6a). This indicates that in the holding period during rehabilitation and post-treatment, regrowth rates are relatively low, vary by facility location, and are still a barrier to recovery for some individual turtles.

**Figure 6.**
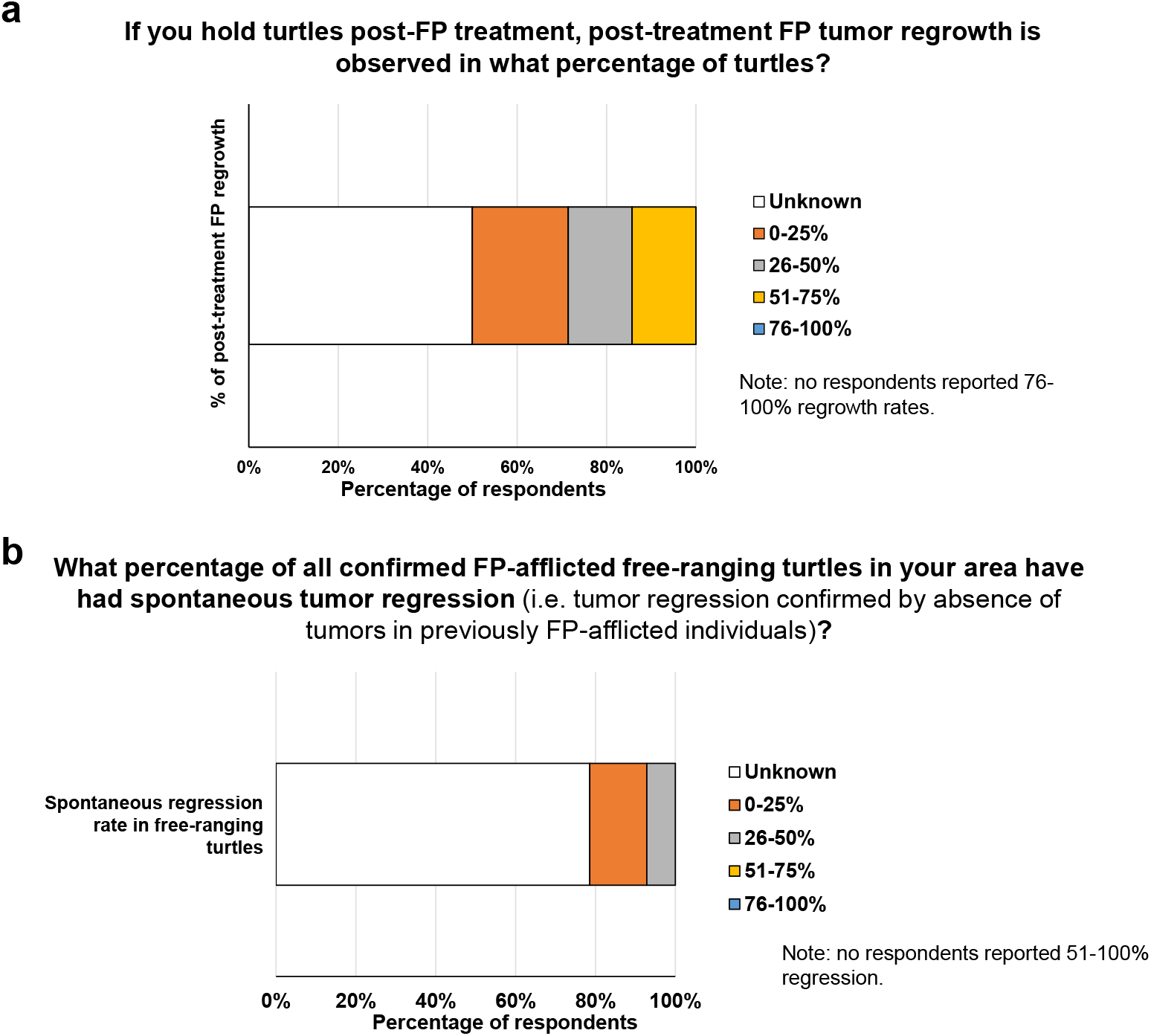
**a)** Percentage of turtles exhibiting post-treatment regrowth of FP tumors. **b)** Percentage of FP-afflicted free-ranging turtles exhibiting spontaneous tumor regression.

The majority of medical respondents (78.6%) reported not knowing the percentage of all confirmed FP-afflicted free-ranging turtles in their area that have had spontaneous tumor regression (i.e., tumor regression confirmed by absence of tumors in previously FP-afflicted individuals, not estimates based solely on tumor phenotype) (Fig. 6b), indicating an important knowledge-gap. For the medical respondents who reported regression in free-ranging FP-afflicted turtles, 14.3% reported a spontaneous regression rate ranging between 0 and 25%, while 7.1 % of respondents reported a spontaneous regression rate in free-ranging turtles of 26-50% (Fig. 6b).

Most medical respondents (92.9%) can identify a turtle that re-strands after previously being released (e.g., from identification tags). Nearly a third of respondents (28.6%) have released FP-free turtles (never had FP or recovered from FP after treatment) that were FP-positive when they later re-stranded, while an identical percentage of respondents (28.6%) never had a released FP-free turtle re-strand with FP tumors. The remaining medical respondents (42.7%) did not know the answer to this question for their facility. 21.4% of medical respondents were aware of previously released sea turtles that were treated for FP and were subsequently documented to have successfully nested. Between two and ten individual turtles were reported by each respondent to be known to have subsequently nested. Of the medical respondents that did not have documented evidence of subsequent nesting, 35.7% reported that FP rehabilitation had not been occurring for long enough in their area for treated FP-afflicted juveniles to mature to nesting age, while 35.7% reported that it was because adequate records of release or nesting were not available. The remaining respondents reported that, in their area, either FP was not present, FP monitoring was not conducted, or FP response strategies were not yet in place (i.e. no FP treatments).

The theme of research (and stakeholder) collaboration was raised in a number of responses. This was mirrored in the response to a direct question on this topic, with 82.9% of respondents considering large-scale multidisciplinary collaboration ‘extremely important’ for advancing the field of FP research, and only 2.4% of respondents consider it of no importance (Supplementary Fig. 1c). Smaller targeted research teams were also considered important, but 2.4 times more respondents reported that collaborative efforts are ‘extremely important’ (Supplementary Fig. 1c).

## Discussion

The last major reviews of the scientific literature on FP were conducted in 2015^15,55^. In the intervening years between those reviews and this survey, FP has spread to new geographic locations, and is now found further from the equator than before^15,55-61^. Simultaneously, FP rates continue to increase in many affected locations^52,55,62-65^. Worryingly, tumor burdens of afflicted individuals and the rate of internal tumor occurrence is also anecdotally on the rise in some locations (personal communication, Bette Zirkelbach and Dr. Brooke Burkhalter). Alarmingly, the results of this survey suggest a further deterioration of the global FP situation, with the majority of respondents (74.4%) considering FP to now be a panzootic (animal pandemic) event. Most respondents report FP prevalence to be increasing in their area, with only 4.7% of respondents seeing a decrease in disease prevalence. Over half of respondents reported that FP tumor burdens on individual turtles are increasing, with only 2.4% reporting decreases in their area. Taken together, FP represents a growing threat to sea turtle survival and conservation, particularly to green turtles, with 62.8% of respondents classifying FP as a strong or major threat. Of all threats listed in the survey, marine pollution, climate change and fisheries bycatch were all ranked as major threats by approximately half of respondents. It is well recognized that sea turtle species face myriad threats to their survival^8,9,66,67^.

In addition to considering FP a panzootic, the vast majority of respondents consider the disease to be cancerous, a progression from earlier thinking on the disease. This aligns with recent transcriptomic and genomic analyses, which have revealed that FP shares many oncological molecular and mutational similarities with human cancers^17,26,41,42^. The similarities with human cancers have allowed human anti-cancer treatments to be applied to sea turtles that are afflicted with FP, thus harnessing readily available drugs and facilitating more effective treatment^41,48,49^.

Basic FP monitoring, organization and infrastructure is lacking in many areas, perhaps due to the continued spread of this disease to new geographic areas. Incidence recording and longitudinal studies were highlighted by respondents as the most beneficial population-level information for FP management. Even in areas with robust monitoring of FP prevalence in stranded turtles, such as Florida, Hawai’i, Texas and Brazil^55,62,63,68,69^, there is a lack of centralized, readily accessible data on in-water FP rates, and a paucity of longitudinal surveillance of progression, relapse and mortality rates of individual turtles.

Given the complex multi-factorial nature and long temporal dynamics of this disease, more research is required to home in on the treatment and mitigation efforts most likely to produce the largest conservation gains. While there was a general consensus among participants that viral, host and environmental factors contribute to the disease, more research is required to elucidate the specific role of the ChHV5 and CmPV1 viruses, and to determine FP’s specific environmental co-trigger(s). Participant responses largely corroborated the current opinions in the scientific literature about FP’s cofactors^2,15,55^. While pollution and eutrophication are correlated to FP prevalence^15^ (scientific literature and survey results), it has not yet been established whether general habitat degradation or sea turtle debilitation are driving factors, or whether it is specific carcinogens, diet changes or immune suppressors that primarily contribute to the FP panzootic. Additionally, seasonality and temperature have been linked to FP tumor growth^15,70^, and 46.5% of respondents considered sea surface temperature to be a driver of FP in their area. However, causal relationships between temperature and FP viral activity or host cancer cell growth rates have not been formally established, although some other classes of viruses are known to be temperature sensitive^71^.

While the survey results illuminate the urgent need for continued and novel FP research, they also highlight that a lack of adequate funding is creating a bottleneck to conducting FP research, which is slowing progress on understanding, combating and mitigating this disease. Other impediments to FP research include onerous regulatory requirements, low recapture rates and limited infrastructure and training. A cross-stakeholder approach from permitting agencies to funders to education providers, conservationists and communities is therefore required to address these bottlenecks and enable FP research to advance more rapidly.

As with FP research, FP rehabilitation was broadly reported to be under-funded with insufficient capacity globally. Given the continued spread of the disease, there is a critical need to train not just more researchers, but also more veterinary and husbandry staff to specifically house and treat FP-afflicted sea turtles. Some respondents highlighted the complete lack of facilities capable of catering to FP-afflicted sea turtles in their areas, and some respondents even reported a complete absence of dedicated general sea turtle rehabilitation facilities. While there is some heterogeneity in capacity, infrastructure, FP medical expertise, and euthanasia decisions, the modes of assessment, diagnosis, surgery and drug treatment for FP were more homogeneous (from respondents in areas with facilities to accommodate FP patients). The survey highlighted knowledge gaps in relation to tumor regrowth rates and post-release outcomes, as well as a paucity of centralized, readily accessible, comprehensive patient progression and outcome data. Gaps in record-keeping of basic disease statistics that can inform research and clinical decision-making likely arise as a consequence of underfunding in the FP rehabilitation sector, and the voluntary *ad hoc* nature of sea turtle rehabilitation, in which each facility has its own practices and data recording priorities.

This survey provides an informative overview of global FP research and FP trends, as well as baseline data which can be built upon. Because the conference was virtual, had an intentionally low financial barrier to participation (i.e., not limited to those with funds to travel), and the survey was conducted online and open to all self-reported FP experts, the survey was equitable and inclusive, capturing FP information and leading opinions from a wide range of geographic areas. While the survey achieved a wide global spread of respondents, and a proportionally high number of respondents, given that sea turtle FP research is still a relatively niche field, there is scope to increase the respondent number and achieve a more global distribution of respondents, especially from emerging disease locations. This survey should be utilized as a basis for a recurring (tri-annual) overview of sea turtle FP status, trends and research. Future surveys can build on this foundation and baseline data, and the survey can potentially be offered in more languages to help facilitate responses from non-English speakers.

The Ocean is the cradle of multicellular life. As such it can teach us much about biological processes, including oncogenesis. Marine species are currently underutilized as natural models of cancer and as sources of novel drug discovery. In addition to marine microbial-based drug discovery, long-lived multi-cellular marine species with wide ranging environmental and pathogen exposures, such as sea turtles, have much to teach us about the processes driving oncological transformation^2,6,72,73^. Sea turtles normally have robust tumor defenses, and understanding these and how they have been overcome in the case of FP will likely prove to be highly informative for human oncology. Such novel insights will not only benefit humans but will have return benefits to wildlife species. For example, host oncogenic signaling occurring in human cancer are mirrored in sea turtle FP^26,74^, and human anti-cancer medications have been shown to be effective for treating sea turtle FP tumors. Any new anti-cancer compounds derived from marine species are likely to have an equal chance of being beneficial for humans, sea turtles and other cancer-afflicted marine species.

## Summary and recommendations

Taken together, the results of this sea turtle FP survey highlight that this panzootic disease poses a serious and worsening conservation challenge. While sea turtles face a worrying array of threats, all of which require action, the nature of FP is different to the other principle threats, which tend to be abiotic. As a multifactorial disease, FP requires a different approach, including development of specific FP-focused expertise and capacity, for research, rehabilitation and mitigation. The scale and severity of the FP threat requires concerted collaborative efforts to tackle the disease, from local stakeholders, responders, and medical professionals, to governmental responses and international training and research initiatives. To effectively tackle this disease, improved financial support for rehabilitation capacity and hospital, lab and field-based research inquiry is required. Major knowledge gaps identified by the respondents that would benefit from increased focus are tumor regrowth, regression and mortality rates, transmission mechanisms and coordinated incidence recording.

Immediate needs highlighted by the survey respondents include improved surveillance of the disease, rapid and equitable data access, advanced training, rehabilitation, research and mitigation resourcing. Short-term goals include establishing care facilities where they are lacking, increasing facility capacity in areas with rising FP prevalence, and research to improve treatment options and causatively identify the viral and environmental triggers. Medium-term goals to improve understanding of the disease causes and consequences should then be translated into the enactment of evidence-based population level mitigation measures. Admirable and dedicated management, rehabilitation and research efforts have occurred over recent decades. However, as the threat of FP spreads, more collaborative and inclusive endeavors will maximize tangible progress in the field and translate advances into measurable gains in sea turtle conservation.

## Acknowledgements

Deepest thanks to all survey respondents for their valuable time, all attendees and presenters at the symposium, and all those who made the running of the symposium possible, especially staff and students of UF Whitney Laboratory. Sincerest thanks to the National Save the Sea Turtle Foundation for enabling and supporting the survey and the associated symposium.

## Figure Legends

**Supplemental Figure 1.**
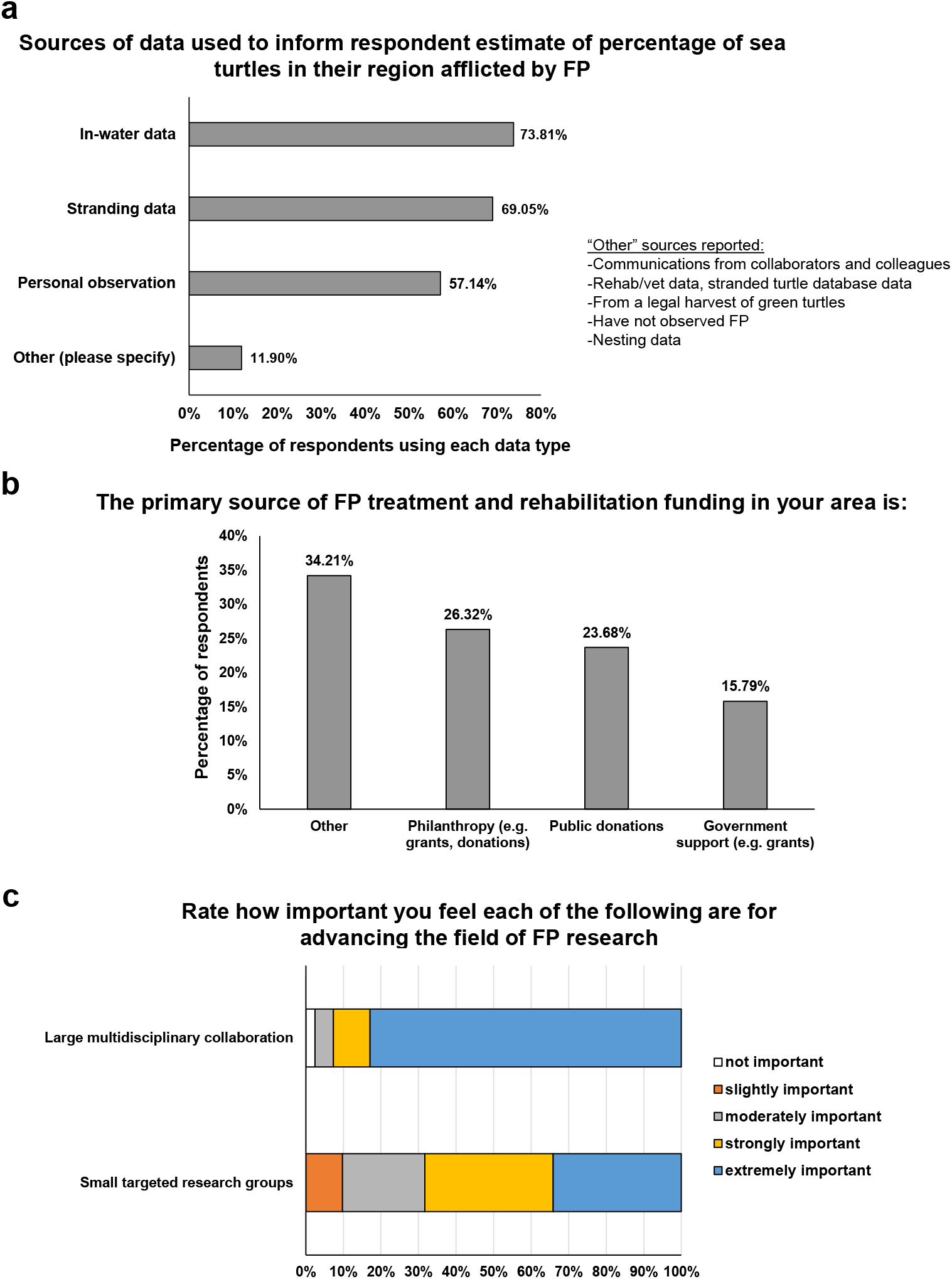
**a)** Sources of data used to inform respondent’s estimates of percentage of sea turtles in their region afflicted by FP. **b)** Primary source of FP treatment and rehabilitation funding in the respondent’s area. **c)** Percentage of respondents that consider large multidisciplinary collaboration or small targeted research groups to be not important, slightly important, moderately important, strongly important, or extremely important for advancing the field of FP research.

